# Properdin oligomers adopt rigid extended conformations supporting function

**DOI:** 10.1101/2020.11.13.381772

**Authors:** Dennis V. Pedersen, Martin Nors Pedersen, Sofia M.M. Mazarakis, Yong Wang, Kresten Lindorff-Larsen, Lise Arleth, Gregers R. Andersen

## Abstract

Properdin stabilizes convertases formed upon activation of the complement cascade within the immune system. The biological activity of properdin depends on the oligomerization state, but whether properdin oligomers are rigid and how their structure links to function remains unknown. We show by combining electron microscopy and solution scattering, that properdin oligomers adopt extended rigid and well-defined conformations that are well approximated by single models of apparent n-fold rotational symmetry with dimensions of 23-36 nm. Properdin monomers are pretzel shaped molecules with limited flexibility. In solution, properdin dimers are curved molecules whereas trimers and tetramers are close to being planar molecules. Structural analysis indicates that simultaneous binding through all binding sites to surface linked convertases is unlikely for properdin trimer and tetramers. We show that multivalency alone is insufficient for full activity in a cell lysis assay. Hence, the observed rigid extended oligomer structure is an integral component of properdin function.

## Introduction

The complement system is an essential aspect of innate immunity providing a first line of defense against invading pathogens as well as maintenance of host homeostasis. The complement cascade is activated when circulating pattern recognition molecules recognizes molecular patterns on a pathogen, dying host cells or immune complexes. Activation can initiate through the classical pathway (CP), the lectin pathway (LP) or the alternative pathway (AP) where the AP also provides an amplification loop for the two other pathways (1). In all three pathways, labile protein complexes known as C3 and C5 convertases are assembled. These convertases conduct proteolytic cleavage of complement component C3 and C5, respectively, resulting in the generation of opsonins (C3b and iC3b), anaphylatoxins (C3a and C5a) and assembly of the membrane attack complex (reviewed in (1)).

In the alternative pathway, the C3 convertase C3bBb is formed when a complex between C3b and the serine protease factor B (FB) is activated by factor D. At a high surface density of C3b, this C3 convertase becomes a C5 convertase. Properdin (FP) is a positive regulator of these convertases. FP is a 53 kDa protein composed of an N-terminal TGF-β binding (TB) domain followed by six thrombospondin type I repeats (TSR1-6). The protein is heavily post-translationally modified carrying one N-linked glycan, four O-linked glycans and 14-17 C-mannosylated tryptophan residues in the WxxW motifs present in TSR1-6 (2,3). FP is produced predominantly by monocytes, T-cells and neutrophils and circulates as oligomers and is primarily found as dimers, trimers and tetramer with a 1:2:1 molar distribution in plasma at a concentration of 5-25 μg/ml (4,5). The functions of FP in relation to AP convertases are well established. 1) FP enhances the recruitment of FB to C3b and thereby stimulates proconvertase assembly; 2) FP slows the dissociation of C3bBb 5-10 fold; 3) FP directly competes with factor I resulting in decreased irreversible degradation of C3b to iC3b. Other suggested functions of FP are as a C3b independent pattern recognition molecule capable of triggering the AP (reviewed in (4)) and as a ligand for the NKp46 receptor on innate lymphoid cells (6). The importance of FP in innate immunity and homeostasis is demonstrated by individuals with FP deficiency (PD). PD is a rare X-linked disorder which can be divided into three subtypes: type-I (complete lack of FP), type-II (1-10% of normal plasma FP level) and type-III (normal plasma level but dysfunctional FP). All PD types are characterized by reduced AP activity resulting in impaired bactericidal activity and increased susceptibility to *Neisseria* infections and sepsis (7).

Classic negative stain (ns) EM studies of FP dimer, trimer and tetramers revealed that FP oligomers contain compact eye shaped vertexes connected by thin connecting structures (8,9). Alcorlo and coworkers presented the first 3D reconstruction of the FP eye shaped vertices in oligomers and 2D classes of the FP-C3bBb convertase complex (10). Recently, crystallographic structures demonstrated that FP oligomers are formed upon interaction of the TB domain and TSR1 from one FP monomer with TSR4, TSR5 and TSR6 from a second FP monomer (Fig 1A). In addition, it was established that the binding site for C3b is formed by FP TSR5 in conjunction with a large loop from TSR6 (2,3,11). Despite prior attempts to analyze FP dimers and trimers with small angle X-ray scattering (SAXS) and analytical ultracentrifugation (12), detailed information regarding the structure and the dynamic properties of FP oligomers is missing. Here we present for the first time a structural description of intact oligomeric FP obtained through a combination of nsEM and SAXS. Based on the recent crystal structures of monomeric FP, we are able to annotate all FP domains in monomeric, dimeric, trimeric and tetrameric FP in EM 2D classes. Pair distance distributions and atomic models based on solution scattering suggest that the FP oligomers are rather rigid in solution despite their very open structure and that their average conformations have cyclic symmetry. In addition, we demonstrate that the defined structure of FP oligomers is crucial for their biological function.

**Figure 1.**
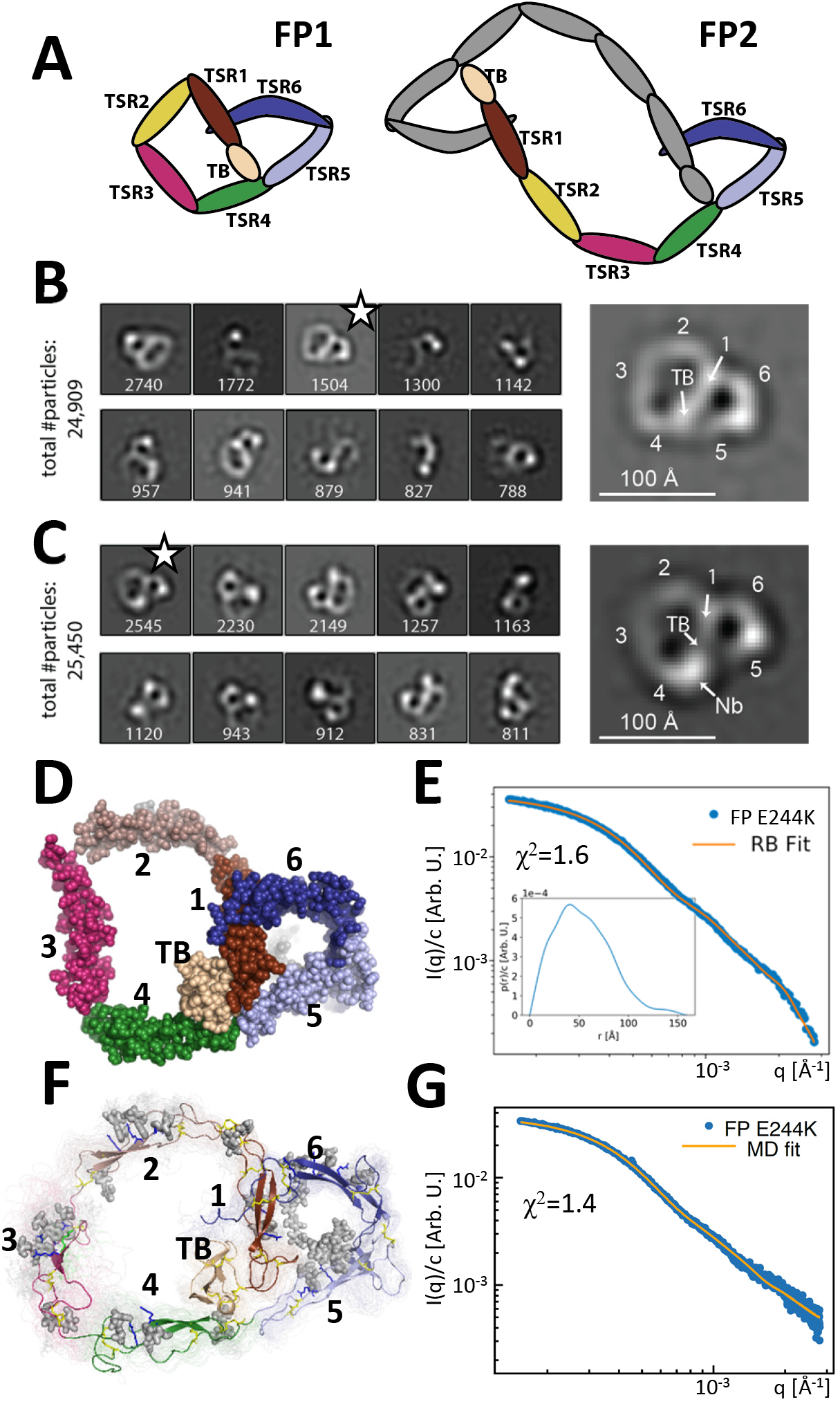
Principles of properdin architecture and the structure of FP1. A) Schematic representation of the FP1 monomer and FP2 as an example of an oligomer, where one subunit is colored grey while the other is colored according to the domain structure as for FP1. B) The 10 most populated nsEM 2D classes obtained with FP1 with the number of particles indicated. A magnified view of the 2D class marked by star is shown to the right. C) As for panel B, but for the FP1-hFPNb1 complex. Compared to FP1, an additional mass marks the location of hFPNb1 and hence TSR4 enabling assignment of the TB domain and the six thrombospondin repeats in the magnified view to the right. D) Representative atomic model of FP1 E244K derived by rigid body modelling against the SAXS data. E) Comparison of SAXS experimental data (11) and the fitted scattering curve corresponding to the FP1 E244K model presented in panel E. Insert to E: *p(r)* function derived from the SAXS data. F) Conformational ensemble of FP1 E244K sampled by a one μs MD simulation represented by 100 frames with 10 ns interval shown as transparent tubes. The starting model is displayed as a cartoon with the glycans and glycosylated residues in grey stick representation. Disulfide bridges are represented by yellow sticks. G) Comparison of SAXS experimental data and the scattering curve obtained from MD ensemble after refinement using the Bayesian maximum entropy approach, the two curves fit with χ^2^ = 1.4. The minor difference in the experimental data apparent at the highest q-values in panels E and G is due to subtraction of a constant by CORAL in panel E.

## Results

### The FP monomer is pretzel shaped

Human FP with a C-terminal His-tag was expressed by HEK293F cells and purified by affinity chromatography. The different FP oligomers were subsequently separated by cation exchange and size exclusion chromatography (SEC). As expected, both recombinant and plasma derived FP eluted in multiple peaks corresponding to the different oligomerization states (Fig S1A-B). Besides dimeric, trimeric and tetrameric FP we also observed a small amount of monomeric FP (FP1) in both recombinant and plasma derived samples eluting after 14.0 ml which is similar to the elution time for the FP E244K variant described previously (11). E244K However, we were able to purify a significant amount of FP1, which allowed us to obtain negative stain EM (nsEM) data for a wild type FP monomer. The resulting 2D classes revealed a flat pretzel shaped molecule with apparent overall projected dimensions of 95×115 Å containing a small and a large ring-shaped structure sharing one edge (Fig 1B). Due to strong preference in the orientation of FP1, we could not obtain a 3D reconstruction of the molecule. Nevertheless, by analyzing nsEM 2D classes obtained for FP1 in complex with the nanobody hFPNb1 (Fig. 1C) and by comparison with the known crystal structures of FP, we could identify the positions of the individual TSRs in the 2D classes of FP1 and nanobody bound FP1. The small ring of the FP1 molecule corresponds to the well described FP “eye” formed by the TB domain, TSR1, TSR5 and TSR6 (2,3), whereas the larger ring is formed by TSR2, TSR3 and TSR4 together with the TB domain and TSR1.

Based on the crystal structure of the recombinant two-chain monomer FPc in which TSR3 and TSR4 are not connected (2,11), we manually constructed an atomic starting model in accordance with the FP1 2D classes and performed rigid body refinement of this model against existing SAXS data obtained for the FP1 carrying the E244K mutation (11). Distance restraints secured appropriate distances across the TSR1-TSR2, TSR3-TSR3 and TSR4-TSR4 interfaces and maintained an appropriate distance between the disulfide bridged Cys132 in TSR1 and Cys170 in TSR2. The resulting models clustered tightly and fitted the data with χ2 = 1.6 (Fig 1D-E and Fig S2B). The SAXS models strongly resembled the EM 2D classes obtained for the wt FP1 monomer. The *p(r)* function derived from our published SAXS data FP E244K monomer (11) approaches zero at about 13 nm with a small tail stretching out to a *D_max_* at 15 nm (see insert to Fig 1E). This agrees with the maximum extent of 14 nm in the nsEM 2D class projections.

To obtain insight into the dynamic properties of FP E244K, we conducted explicit solvent atomistic molecular dynamics simulations of a complete model of FP E244K at microsecond timescale initiating from our SAXS model obtained by rigid body modeling (Movie S1). The model included the Asn-linked complex glycan, Ser/Thr O-linked glucose-fucose, and the C-linked mannosylations on TSR tryptophans (Fig 1F). The SAXS curve calculated from the ensemble of conformations sampled by molecular dynamics simulations is in good agreement with the experimental SAXS data (χ^2^ = 2.2). By using experimental data to guide the refinement of the conformational ensemble, the fit could be further improved to be χ^2^ = 1.4 using the Bayesian Maximum Entropy (BME) method (Fig 1G). Models in this ensemble resembled the FP E244K output models from the SAXS rigid body refinement (Fig 1D) and the 2D classes obtained by EM of wild type FP1 (Fig 1B). Although the mutation E244K is expected to weaken the Trp-Arg stack in TSR3, throughout the MD simulation the Trp-Arg stack remained intact (Movie S2). However, two independent MD simulations together suggested that TSR3 inserted between TSR2 and TSR4 is capable of rotating as a rigid-body with limited conformational flexibility within domain (Fig S3C-F). This domain rotation was apparently facilitated by high mobility of the TSR4 loop region Asn285-Phe295 facing TSR3 which is in excellent agreement with weak electron density we observed for this region in two crystal structures (2). At the TSR2-TSR3 junction, the corresponding TSR3 loop Ser225-Pro237 facing TSR2 likewise exhibited flexibility (Fig S3C-D and Movie S1). For the remaining parts of FP E244K, the simulation did not indicate significant mobility, and in particular the two loops in TSR5 and TSR6 involved in convertase recognition (2,3) did not exhibit significant changes in backbone conformation (Fig S3C and Movie S1). Overall, the nsEM 2D classes for wt FP1 and the SAXS data for FP E244K provided the first quasi-atomic experimental model for the monomeric FP1 and FP E244K. Although FP1 like FP E244K probably have limited biological activity, it is present in plasma FP at very low concentrations (Fig S1B).

### Pseudosymmetric FP oligomer conformations are observed by EM

We next analyzed the FP dimer (FP2) isolated using SEC by negative stain EM. Of the 51% of particles that were classified, 8/10 appeared in similar 2D classes suggesting that a limited number of related elliptical dimer conformations are present on the EM grids (Fig 2A). The overall projected dimension of FP2 in a typical 2D class was 27×13 nm with an inner opening of 11×5 nm. However, in some classes the central opening appeared broader and the molecule less extended. At each end of the dimer, the characteristic eye shaped structure was present and the TSRs could be assigned by comparison with the 2D classes of FP1. Interestingly, TSR3 of FP2 appeared to connect TSR2 and TSR4 in an almost linear arrangement (Fig 2A) which is in contrast to the kinked conformation observed for FP1. The 2D classes suggested that the two monomers in FP2 are related by an approximate two-fold rotation axis (C_2_ symmetry) perpendicular to the plane of the dimer.

**Figure 2.**
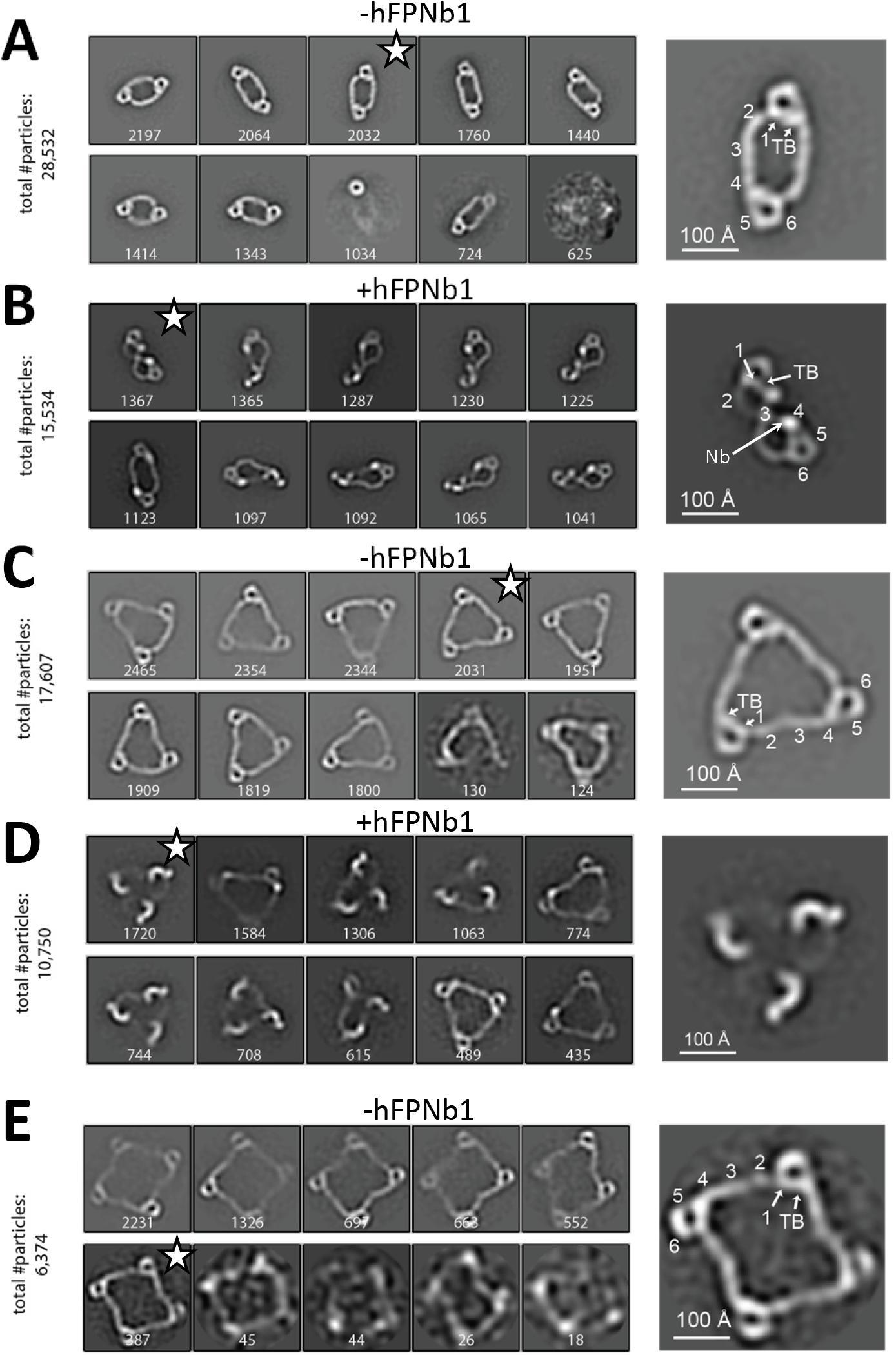
Negative stain EM analysis of FP oligomers. For all cases, the 10 most populated nsEM 2D classes obtained are displayed with the number of particles indicated. A). The 2D classes obtained with FP2 and a magnified view of the 2D class marked by star is shown to the right with the putative domain assignment indicated for one of the two monomers in the dimer. Notice the difference in curvature at TSR2 and TSR4 of the connecting arms, which facilitates the domain assignment. B) As panel A, but with the FP2-hFPNb1 complex. The double pretzel conformation is present in two classes, while others feature the elliptical shape or intermediates. C) The 2D classes obtained with FP3 reveal a flat molecule structure of apparent C3 symmetry. D) The FP3-hFPNb1 complex. The 2D class to the left reveals an FP3 conformation with an apparent C3 symmetry that is radically different from the cyclic appearance of FP3 in panel C, but 2D classes presenting flat cyclic FP3 and intermediates are also present. E) The 2D classes obtained with FP4 suggest a planar extended molecule with apparent C4 symmetry.

Whereas the FP2 2D classes above concurred with prior EM studies (8,10), radically different 2D classes were also obtained when hFPNb1 was included (Fig 2B). In one extreme 2D class, FP2 appeared as a folded dimer adopting a double-pretzel structure with a maximum dimension of 21 nm where the two connecting arms are crossing over at TSR3, and like in FP1, the TSR2-TSR3 angle is sharply bend (Fig 2B). This FP2 conformation appeared to have a C_2_ symmetry axis lying in the plane of the molecule in contrast to unbound FP2 where the apparent C_2_ axis was perpendicular to the plane of the molecule. Other 2D classes obtained with hFPNb1 presented conformations that were intermediate between the open dimer observed in the absence of hFPNb1 and the folded dimer. A comparison of the SEC profiles supported significant conformational effects of hFPNb1 on FP2 in solution prior to EM sample preparation (Fig S1D). Importantly, hFPNb1 does not influence the oligomer distribution as shown previously (13).

We next analyzed the trimer FP3 by nsEM, where eight very similar 2D classes contained >90% of the particles picked (Fig 2C). In these classes, FP3 appeared in projection as flat triangular molecules with pseudo C_3_ symmetry having maximum edges of 28 nm with mainly straight connections formed by TSR2, TSR3 and TSR4 between the vertices. Again, when FPNb1 was present, some 2D classes contained a less planar FP3 with a maximal dimension of 24 nm where only the eye shaped vertices could be unambiguously identified (Fig 2D). Other FP3-hFPNb1 2D classes featured molecules that were flat and circular like those obtained in the absence of hFPNb1 or intermediate between these two extremes. Finally, we analyzed the tetramer FP4 by nsEM and again >90% of the picked particles contributed to 2D classes with flat molecules of pseudo C_4_ symmetry with a maximum extent of 38 nm (Fig 2E). For both FP3 and FP4 2D classes, a kink was occasionally observed, presumably at the TSR2-TSR3 interface, in at least one of the TSR2-TSR3-TSR4 connections. In summary, our nsEM analysis revealed unambiguous 2D classes for all the naturally occurring FP oligomers, and in all three cases the vast majority of classified particles were present in rather similar classes representing flat molecules with apparent C_n_ symmetry. Furthermore, the oligomer conformations observed in our 2D classes were not rare since the sum of particles used for the 2D classes for all three oligomers represented at least 50% of the particles picked (Fig. 2). Intriguingly, we also observed that the TSR4 specific hFPNb1 could induce alternative folded conformations of the TSR2-TSR3-TSR4 arms in FP2 and FP3 that appeared to be in equilibrium through intermediates with their elliptical and flat triangular conformations.

### The solution conformation of FP oligomers

Taking an orthogonal approach to our nsEM analysis of the FP oligomers, we collected SEC SAXS and static SAXS data for FP2, FP3 and FP4 that was SEC fractionated prior to SAXS analysis to obtain samples optimized with respect to the relevant FP oligomer (Figs 3A-C and S2A-G). It is noteworthy that the SAXS data of all three oligomers exhibit characteristic bumps which are also reflected in their *p(r)*-functions (see inserts to SAXS data and fig S2H). The presence of these pronounced features clearly indicate that the oligomers are rather rigid and well-defined. If conformational freedom had been present, this would smear out the SAXS data. The *p(r)*-functions of FP1, FP2, FP3 and FP4 all exhibit an initial bump at around 4 nm corresponding well to the repeated distance across the eye that is also visible in the nsEM pictures of all three oligomers (see also fig S2H). The FP2 has a *D_max_* value of 23 nm. Along with a second well-defined peak at around 8 nm, which is most likely related to the distance between the two antiparallel arms connecting the two eyes, the *D_max_* value indicates that the FP2 solution structure is in better agreement with the extended conformation of FP2 (Fig 2A), than it is with the more compact twisted conformation induced by hFPNb1 (Fig 2B). Finally, the *p(r)* of FP2 has a broad peak at 16.5 nm corresponding well to the distance between the two eyes. The *D_max_* of the FP3 and FP4 *p(r)* functions are, respectively, 25 and 36 nm (Fig 3B-C and Fig S2H) and in good agreement with their larger sizes also seen in the nsEM 2D classes (Figs 2C and 2E). For the FP3 and FP4, the *p(r)* middle peak at around 8-10 nm appears broader, less pronounced and moves to higher values as the oligomer size increases. FP3 and FP4 have a high *p(r)* peak at 18 and 19.5 nm, respectively, which is most likely the result of neighbor eye-eye distances. The increase of the eye-eye peak position when comparing FP2, FP3 and FP4 suggests that the eyes reorient towards a more planar structure as the oligomeric state increases. Furthermore, the FP4 *p(r)* function exhibits an additional small shoulder at 28 nm which corresponds well to the less frequent diagonal eye-eye distances. For a quadratic structure, as suggested by the nsEM, these would appear at 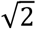 times the position of the neighbor eye-eye distances at 18-19.5 nm as they indeed do.

**Figure 3.**
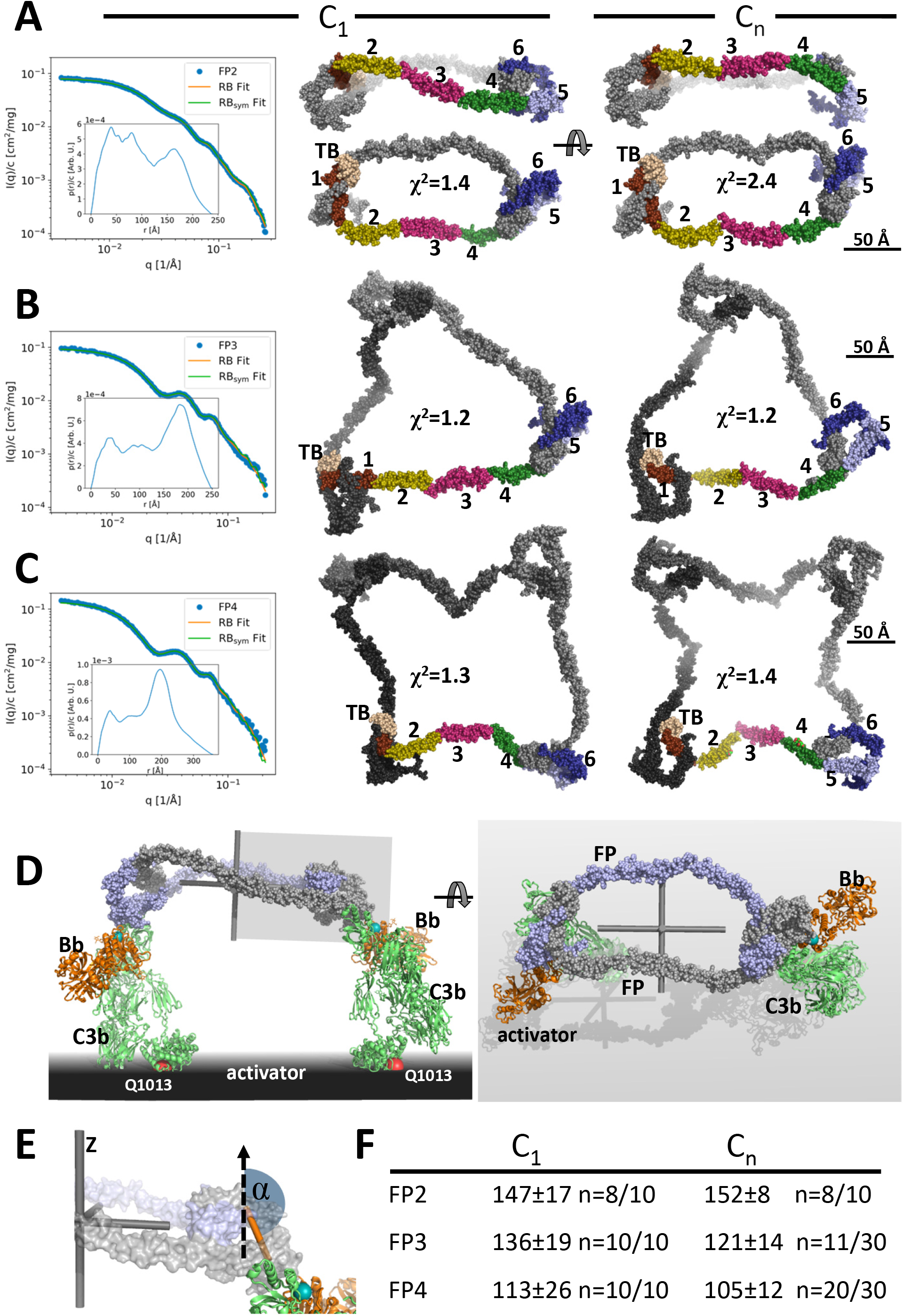
Models of FP oligomers obtained by SAXS rigid body refinement. A-C) Left: SAXS data (blue) and model fits (orange) corresponding to FP2, FP3 and FP4 structures obtained with C_1_ symmetry. Inserts plot the corresponding *p(r)* functions. Middle: Models fitted using C_1_ symmetry. Right: Models fitted assuming Cn symmetry. The C_1_ models were chosen as the most representative model in the largest cluster from the 10 generated, whereas the Cn models were those amongst 10 or 30 models that according to their χ^2^ value were in best agreement with the experimental data, see panel F. In panel A, two orientations are displayed to illustrate the curvature of FP2 models. D) Hypothetical model of FP2 from panel A bound to two C3 convertases to illustrate that such a complex could be formed on an activator to which the two C3b molecules (green) carrying the Bb protease (orange) are bound through their Gln1013 (red sphere). The two FP monomers are colored light blue and grey, respectively. To the left is presented a side view and to the right a top view. The principal axes of FP2 are displayed as a grey Cartesian coordinate system. E) Enlarged view of the area outlined with a grey box in panel D. The vertical dashed vector is parallel to the smallest principal axis (labelled “z”) of the FP2 molecule, the orange stick represents the vector in the plane of the FP eye used for calculation of α, the angle between the two vectors. F) The average α values calculated from the SAXS rigid body models. The number of models used for statistics from a pool of 10 or 30 models is indicated by the number n, remaining models were discarded as they had χ2 values that were significantly higher than the n well fitting models.

Using the same strategy and restraints as for FP1, we obtained rigid body models of FP2 in the presence of C_1_ (no) symmetry or a C_2_ symmetry axis with χ^2^ in the range 1.4-3.0 for C_1_ symmetry whereas models with C_2_ symmetry had χ^2^ of 2.4-5.5 (Fig 3A and Fig S2C). The resulting SAXS-based FP2 models all featured an extended FP2 conformation rather than the double-pretzel FP2 observed by nsEM, even if refinement was started from a double-pretzel conformation mimicking that presented in figure 2B. Interestingly, the FP2 SAXS models appeared more curved as compared to the extended conformations present in EM 2D classes. In contrast to the nsEM 2D classes, the rigid body models can not be projected such that the two eye-shapes at the opposite ends of FP2 become visible simultaneously. Hence, either rigid body modelling could not reach the conformation observed in nsEM, or the FP2 conformation is influenced by the stain and contacts with the grid and therefore become excessively flat in our 2D nsEM classes. An effect of the grid is in accordance with a lower solution D_max_ compared to the maximum extent of FP2 in nsEM 2D classes.

Using the same strategy we generated models of FP3 and FP4 by rigid body modelling. In contrast to FP2, the χ^2^ values for the fit of the best models to the experimental data were comparable and in the range 1.1-1.4 indicating that the experimental data could be fitted well with single models. Importantly, the χ^2^ values of the output models generated with C_3_/C_4_ symmetry were similar to those generated with C_1_ symmetry demonstrating that the fit to the data was independent of symmetry (Fig 3B-C and S2D-E). All models generated with C_1_ symmetry were qualitatively similar extended circularized structures with their 3 or 4 eye shaped structures joined by peripheral TSR2-TSR3-TSR4 connecting arms in agreement with the FP3 and FP4 nsEM 2D classes. As for FP2, in the models generated with C_1_ symmetry, it was never possible to obtain projections of these SAXS models in which all eyes in FP3 and FP4 were visible. Otherwise the SAXS models had strong resemblance to the 2D classes including an occasional kink in a TSR2-TSR3-TSR4 connection (Fig 3B-C). When C_3_/C_4_ symmetry was assumed during refinement, some models were trapped in a local minimum and gave rise to highly elevated χ^2^ values and models that were distinctly different from those with the lowest χ^2^ values. In contrast, the best fitting models were all open and extended with TSR2-TSR3 at the periphery similar to those obtained with C_1_ symmetry. For some of these models, it was possible to visualize all eyes simultaneously in projections (Fig 3B-C) which is in agreement with FP3 and FP4 nsEM 2D classes. Overall, our SAXS analysis of FP oligomers were in agreement with the corresponding nsEM 2D classes and suggests that the average solution conformation of FP3 and FP4 - as expressed in the SAXS data - are well approximated by single models with C3 and C4 symmetry (S2D-E). With respect to FP2, the situation is less clear as larger differences were observed for rigid body modelling with C_1_ and C_2_ symmetry and the resulting models appear more curved than those appearing in the nsEM classes.

### The oligomerization interfaces can undergo very slow exchange

The oligomer distribution of FP in plasma is believed to be stable after secretion, but whether monomer-monomer interactions occasionally loosen up and whether monomers may exchange between oligomers has never been addressed. Our prior purification of the two-chain monomeric FP molecules FPc and FPhtΔ3 (2,11) never suggested that the two chains dissociated under native conditions. However, in size exclusion chromatography performed at low pH it was possible to separate the two FP chains (Fig. S4A) as previously demonstrated for FP oligomers (5). To investigate the stability of the FP oligomerization interfaces, we mixed our two-chain monomer FPc with the two-chain deletion mutant FPhtΔ3 monomer lacking TSR3 (Fig S4B). After incubation at 37 °C, we purified FP molecules containing the His-tagged TSR4-6 tail fragment of FPhtΔ3. Using SDS-PAGE analysis, we observed that over time an increasing amount of the longer head fragment from FPc (TB-TSR1-3) co-purified with the His-tagged tail fragment from FPhtΔ3 while the amount of co-purified short FPhtΔ3 head fragment (TB-TSR1-2) decreased (Fig. S4C). Exchange between the two FP monomers was evident after 30 min and reached equilibrium after 12 hours. To examine monomer exchange into an FP oligomer, we conducted the same experiment with FP2 and FPhtΔ3. We observed that over time, an increasing amount of full length FP from FP2 co-purified with the FPhtΔ3 his-tagged TSR4-6 fragment, and in this case exchange was evident after 2 hours and complete in 6-12 hours (Fig. S4D). In conclusion, these experiments for the first time demonstrated that the oligomerization interfaces in FP can open temporarily and even exchange monomers with a different FP molecule under physiologically relevant experimental conditions. However, the exchange occurs rather slowly in our pure system and is probably not a significant reaction in the extracellular environment after secretion. These results add further support to the concept of stable FP oligomer conformations.

### Convertase binding to FP oligomers

A major outstanding question concerning FP biology is whether the strong correlation between biological activity and oligomer size is partly due to simultaneous binding to multiple C3b molecules and convertases deposited on an activator. In FPn, there are n binding sites for C3b each located at the concave face of TSR5, and our prior structures of FP-bound C3bBb revealed that C3b binds with its major axis roughly parallel to the plane of the FP eye formed by the TB domain, TSR1, TSR5 and TSR6 (14). Hence, as illustrated in Fig 3D for FP2, an oligomer with its n convertase binding sites pointing in the same direction will have the optimal architecture for binding simultaneously to n C3b molecules deposited on an activator. Due to the overall flat shape of FP oligomers, the minor principal axis is the C_n_ rotation axis perpendicular to the plane of the models generated with C_n_ symmetry and a pseudo n-fold rotation axis for models generated with C_1_ symmetry (Fig 3E). We could therefore quantitate the relative orientation of the convertase binding sites in our FP SAXS models by measuring the angle α between an appropriate vector lying in the plane of each FP eye and the smallest principal axis of the oligomer (Fig 3E). At the maximum value, α=180°, the n convertase binding sites in an FP oligomer will point in the same direction and simultaneously binding at all convertase binding sites by C3b on a planar activator appears possible. At its minimum value, α=90°, the convertase binding sites are parallel to the plane of the FP oligomer defined by the major and intermediate principal axes and simultaneous binding to more than two C3b molecules on a planar activator appears unlikely. We observe a clear decrease in α as a function of oligomer size with α~150°, 128° and 109° for our SAXS-based models of FP2, FP3 and FP4, respectively (Fig 3F). A decrease in α with increasing multiplicity is in agreement with the shift of a major peak in the pair distance functions (Fig S2H) described above. This peak largely reflects the separation of neighboring FP eyes, and in curved oligomers this separation will be smaller than in planar oligomers.

### Oligomerization alone cannot rescue FP activity

The activity of FP oligomers in assays exploring complement-dependent erythrocyte lysis follows the order FP4>FP3> FP2 (5), and the two chain monomer FPc and E244K FP1 are also much less active in convertase stabilization on erythrocytes and bactericidal activity compared to oligomeric FP (11). To investigate the importance of FP oligomerization and structure for activity we linked 2, 3 or 4 copies of hFPNb1 with glycine-serine linkers and showed that noncovalent FPc oligomers of increasing molecular weight linked by the multivalent hFPNb1 molecules could be formed (Fig 4A-B). Next, we compared the activity in erythrocyte lysis in FP deficient serum of free FPc and nanobody linked FPc oligomers to an FP pool containing roughly equal amounts of FP2 and FP3 (Fig S1F). As expected, the two-chain FPc monomer required a 100 fold higher concentration compared with the FP2/FP3 oligomer pool to elicit a similar degree of lysis (Fig 4C). The activity of the nanobody-linked FPc oligomers were in between the activities of the FP2/FP3 pool and FPc and increased with the hFPNb1 valency in the oligomers. The erythrocyte lysis activity of these hFPNb1 linked FPc oligomers also correlated well with their increasingly slower dissociation from a biotin-C3b sensor (Fig 4D). In conclusion, FP activity can only be partially restored by linking FPc monomers through flexible linkers despite that these nanobody-linked monomers exhibited increasingly stronger binding to a C3b coated sensor compared to the FPc monomer. Hence, the well-defined extended structure of FP oligomers demonstrated by our SAXS and EM data contributes significantly to the biological activity of FP oligomers.

**Figure 4.**
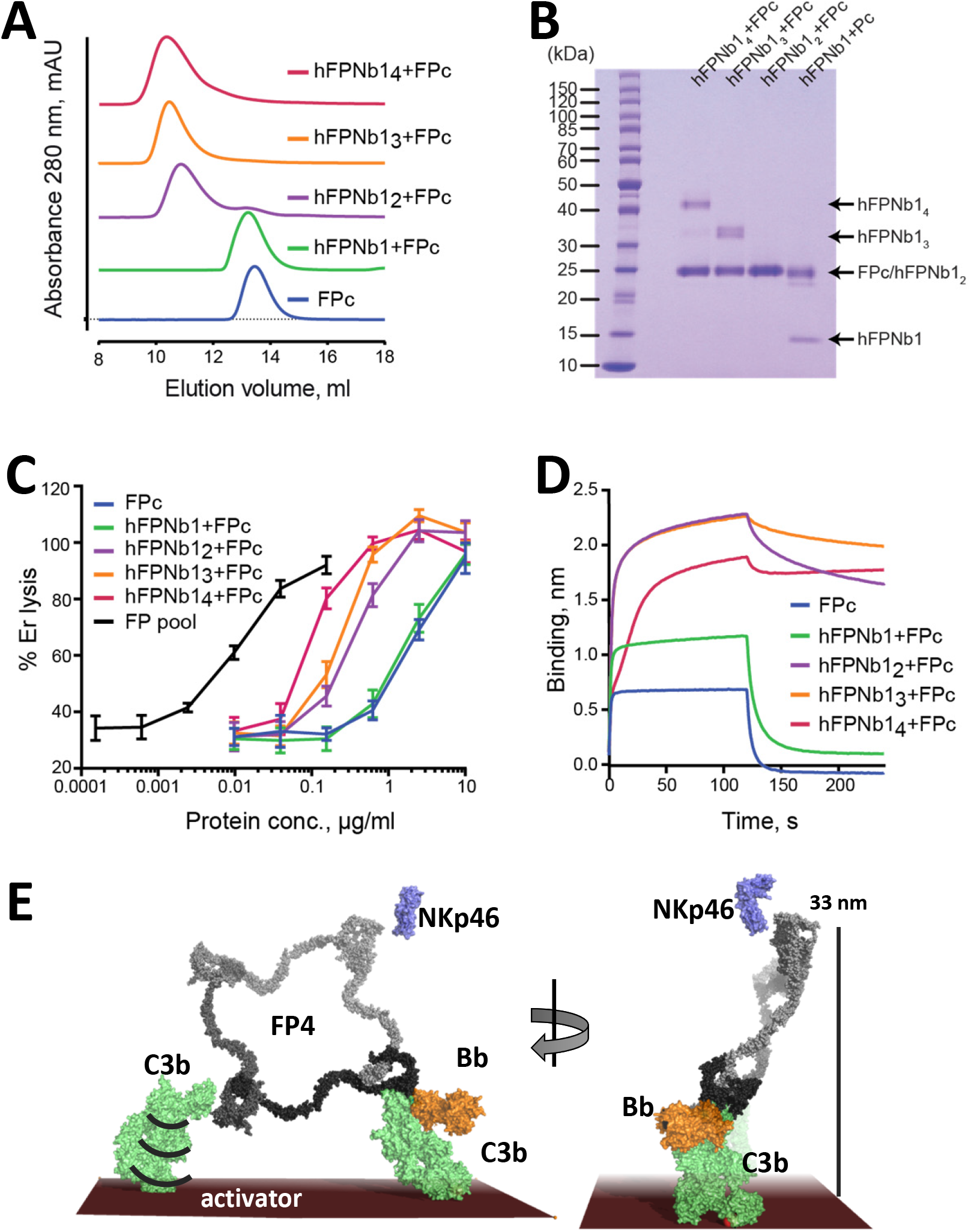
The biological activity of nanobody linked FP oligomers is lower than that of an FP2/FP3 oligomer pool. A) SEC profiles showing that hFPNb1 oligomers form stable oligomers with FPc. B) SDS-PAGE analysis of the peak fraction from each SEC experiments in panel A confirming that all hFPNb1 variants form stable complexes with FPc. C) Assay where lysis of rabbit erythrocytes (% of lysis in pure water) in FP depleted serum is used as a measure of AP activity. Erythrocyte lysis is shown as a function of protein concentration. The ability to induce erythrocyte lysis clearly increases with increasing hFPNb1 mediated oligomerization of FPc. Each point corresponds to the average of two independent experiments, both performed in technical duplicates. The error bars represent the standard deviation between the two independent experiments. D) Biolayer interferometry sensorgrams showing the binding of FPc (blue) or preformed FPc-hFPNb1 monomers (green), dimers (purple), trimers (orange), or tetramers (red) to a C3b coated sensor. The sensorgrams demonstrate that binding of hFPNb1 mediated FPc oligomers to C3b is stronger than for the monomeric FPc or the FPc-hFPNb1 complex. In particular, the dissociation rate is observed to be strongly dependent on hFPNb1 valency. E) Model of FP structure-functions relationships using FP4 as example. To the left, one convertase binding site is occupied, a neighboring binding site may form contact with a nearby C3b, proconvertase or convertase. Flexibility of C3b due to the single bond attachment to the activator may favor FP interaction with the second C3b molecule. Right, FP convertase binding sites not pointing towards the activator surface reach >30 nm into solution and appear as ideal for bridging C3b opsonized pathogens with NKp46 presenting innate lymphoid cells. Notice that the NKp46 binding site on FP is presently unknown.

## Discussion

Our demonstration of well-ordered EM 2D-classes and our ability to explain solution scattering data with single models make us suggest that FP oligomers adopt a limited number of fairly stable and overall similar conformations in solution. This is a surprising finding considering that EM 2D classes of an FP oligomer have not previously been presented and the large flexibility at especially the TSR2-TSR3 and TSR3-TSR4 connections required to reach the very tight FP1 conformation (Fig 1B-D) as compared to FP crystal structures (2,3). The rigidity of FP oligomers is directly manifested by the pronounced oscillations in the SAXS data leading to well-defined peaks in the pair distance distributions (Fig 3A-C and S2H) that present sharp peaks between 15-21 nm originating from the separation of neighboring FP eyes. If the oligomers were flexible, these oscillations would be less well defined.

The solution structures of FP2 and FP3 were earlier investigated by Sun and coworkers with SAXS and analytical ultracentrifugation (12) and gave R_g_ values comparable to those presented in Fig. S2A. Their best fitting FP2 models bear weak overall resemblance to our FP2 models, but their best fitting FP3 model is rather different from the models we obtain. Furthermore, an extended model resembling our models of FP3 did not fit the scattering data in the study by Sun *et al*. One reason for these discrepancies is beyond doubt that Sun *et al* did not have a detailed model of FP to base their rigid body modelling on, and in particular they missed the crucial information regarding the stable eye structure formed by the TB domain, TSR1, TSR5 and TSR6 which we used as a single rigid body. They also used a TSR based homology model as a proxy for the N-terminal region that is a TB domain. In addition, the resolution of SAXS data in (12) was much lower with significant noise at q values above 1 nm^-1^ whereas we had only limited noise in the data in the range q<2.7 nm^-1^ used for rigid body refinement.

The binding sites for C3b is formed by FP TSR5 and TSR6 located in the FP eye and oligomerization is not strictly required for C3b binding, stimulation of C3bB assembly, inhibition of C3bBb dissociation and competition with FI. These activities are supported to some degree by the recombinant monomeric two-chain FPc molecule in which TSR2 and TSR4 are not connected by TSR3 (2,11). Furthermore, EM analysis of erythrocytes activated through the classic pathway show tight clusters of 10-40 C3b molecules extending over 40-80 nm (15), which appears well compatible with the separation of C3b/convertase binding sites observed in our structural models (Fig 3A-C).

Our demonstration that FP oligomers are structurally well ordered rather than dynamic modular structures (Figures 2–3) and our finding that biological activity cannot be rescued solely by linking of multiple monomeric FP molecules (Fig 4D) lead us to present a model in figure 4E for the biological function of FP oligomers. Initial monovalent binding occurs when a single FP eye engages with C3b, proconvertase or convertase. Subsequent multivalent binding is likely to be stimulated by FB binding (16) and appears feasible for neighboring convertase binding sites in all three types of oligomers and especially FP2, where the two sites are arranged favorable with respect to each other for binding parallel C3b molecules separated with an average separation of 20 nm (Fig 3D). The higher activity of FP oligomers could derive from; i) An FP oligomer simultaneously binding multiple activator-bound C3b molecules; ii) a high local concentration of C3b binding sites favoring FP rebinding after dissociation; iii) A defined 3D structure of the oligomers e.g. matching the average distribution of deposited C3b. Our prior SPR data showed a 450× lower apparent K_D_ value for immobilized C3b and fluid phase oligomeric FP as compared to the reverse geometry in accordance with contributions from multivalent binding and a high local concentration (11). In contrast, classic binding experiments measuring association to erythrocyte bound C3bB and zymosan-C3b complexes suggested that FP oligomers bind to a C3b opsonized activator in a monovalent fashion (16,17) arguing that the local high concentration of unbound convertase binding sites for monovalent FP-convertase interactions underlies the correlation between biological activity and oligomer stoichiometry. Our cell lysis experiments demonstrate that multivalency alone is not sufficient obtain full biological activity suggesting that the defined 3D structure of the oligomers also contributes significantly to the correlation between activity and oligomer size. Although we do not know in details the oligomer distribution of FP in non-mammals, the domain structure with the TB domain followed by six thrombospondin repeats is with certainty also present in properdin sequences from amphibians, reptiles, birds, teleosts and the agnatha *Petromyzon marinus*. This evolutionary conservation supports that a defined spatial separation of convertase binding sites in FP oligomers is important for its function. For antibodies, it is known that the flexibility of the Fab-Fc hinge enables “walking” over the antigen in search for bivalent attachment on spatially defined multivalent epitopes (18,19). But since our results indicate that FP oligomers are quite rigid, these multivalent molecules seem not to be designed for “walking” over the C3b opsonized activator. Instead, the inherent flexibility of C3b and its attachment through a single covalent bond to the activator may favor multivalent FP-convertase complexes or fast rebinding through a neighboring convertase binding site (Fig 4E).

One reservation with respect to the model of FP structure-function relationships presented in Fig 4E is that our EM 2D classes obtained with the TSR4 binding hFPNb1 nanobody revealed intricate folded conformations and intermediates between the extended flat cyclic conformation observed in the absence of the nanobody (Fig 2B and 2D). This emphasizes, that the FP oligomer structures we present in figure 3 reflect their solution conformation, but the structure may be quite different for oligomers bound to surface bound C3b and convertases on activators and NKp46 on innate lymphoid cells. The variable internal structure of TSR4 and rather different orientations of TSR4 relative to TSR5 observed in available crystal structures (2,3) and the MD simulations of FP1 presented here agrees with the ability of a TSR4 binder to induce profound conformational changes in FP oligomers. TSR4 is not directly part of the convertase binding site, but it is linked to TSR5 that harbors the majority of the convertase binding site. In an FP oligomer, a conformational signal may be relayed from TSR4 to TSR5 upon convertase binding and propagate into the rest of the oligomer. The strong preference of FP oligomers for orienting with the plane of the molecule parallel to the grid suggests that achieving high-resolution cryo-EM 3D reconstructions of these unique extended structures will be an extremely challenging task. Alternatively, FP oligomers bound to a C3b or convertase coated activator model may be studied with cryo electron tomography as pioneered by Sharp and Gros for large assemblies of complement proteins (20,21).

Interestingly, our structures also predict that empty C3b/convertase binding sites in FP3 and FP4 point in a direction opposite to the occupied site(s) due to their α angle being far from 180° (Fig 3F and 4E). We estimate that such unoccupied convertase binding sites in FP3 and FP4 may protrude more than 30 nm from the thioester-activator linkage of the C3b to which an existing monovalent interaction occurs (Fig 4E). Such empty binding sites appear also to be suitable for bridging a C3b opsonized cell with a second cell. The role of FP mediated agglutination – where C3b tagged bacteria lump together – in bactericidal activity is not well investigated. Historically, FP driven agglutination of erythrocytes was associated with large non-physiologically FP oligomers (22). Perhaps more relevant for *in vivo* activity, FP can also bridge C3b opsonized bacteria with host innate lymphoid cells presenting the NKp46 receptor, an interaction shown to be required for survival in an animal model of *Neisseria meningitides* infection (6). Possibly lack of this cell-bridging function is an important element in the phenotypes of FP deficiencies in addition to compromised convertase stabilization due to weaker binding to C3b on opsonized pathogens. Interestingly, administration of recombinant FP with a high content of FP tetramers and higher oligomers was protective in mouse models of infection with *N. menigitidis* and *S. pneumonia* (23). Possibly, the bridging of bacteria and innate lymphoid cells by larger FP oligomers contributed significantly to the beneficial effects of recombinant FP administration. Very recently FP4, but not FP3 and FP2, was shown to act as a C3b independent pattern recognition molecules on bacteria binding soluble collectin 12 (24) suggesting a specialized function of FP4.

Our identification of FP oligomers as rigid molecules with a potential for bridging C3b opsonized cells with other cells may assist future studies analyzing the role of FP in complement driven pathogenesis and facilitate new strategies for therapeutic modulation of FP activity. Inhibition of FP by function-blocking anti-mouse FP mAbs or FP gene deletion has demonstrated beneficial effects in murine models of arthritis (25), renal *ischemia-reperfusion injury* (*IRI*) (26), allergen-induced airway inflammation (27), abdominal aortic aneurysm (28) and atypical hemolytic uremic syndrome (29). Finally, our results should promote experiments clarifying how FP oligomers may act as C3b independent pattern recognition molecules capable of initiating the alternative pathway.

## Acknowledgements, funding sources and conflict of interest

We thank the staff at the P12 beamline at PETRAIII for help during data collection and Karen Margrethe Nielsen and Anette Hansen for technical support. We acknowledge access to computational resources from the Danish National Supercomputer for Life Sciences (Computerome) and the ROBUST Resource for Biomolecular Simulations (supported by the Novo Nordisk Foundation). This work was supported by the Lundbeck Foundation (BRAINSTRUC, grant no. R155-2015-2666) and the Novo Nordisk Foundation (NNF16OC0022058). The authors declare no conflicts of interest in relation to this manuscript.

## Materials and methods

### Protein production and SEC assays

DNA encoding FP with C-terminal TEV-His sequence was generated by site directed mutagenesis and was expressed by transient expression in HEK293F cells as described in (11). FP was purified from cell supernatants using HisExcel column (GE Healthcare) and 1 ml Mono S column (GE Healthcare) as described (14). Fractions containing FP1, FP2, FP3 or FP4 from the Mono S column were pooled and further purified by SEC performed on a 24 mL Superdex 200 increase column (GE Healthcare) at 4 °C with a flow rate of 0.25 mL/min in a buffer containing 20 mM HEPES, 150 mM NaCl pH 7.5. The monomeric FP variants FPc and FPthΔ3 used for exchange experiments were expressed and purified as described in (2,11), respectively. hFPNb1 were expressed and purified as described in (30). SEC analysis of FPc-hFPNb1 complexes were performed with 200 μL samples containing 14 μg FPc in complex with hFPNb1 or its multivalent derivatives in a 4-fold molar excess with respect to the number of hFPNb1 subunits. Samples were incubated in SEC buffer at room temperature for 15 min before injection.

### SAXS data acquisition, analysis and rigid body analysis

SAXS data was collected at the EMBL beamline P12 at PETRA III in Hamburg, Germany (31). The temperature for the sample changer and exposure unit was set to 8°C, and the detector and X-ray energy was configured to give a *q*-range of 0.023 to 7.332 nm^-1^, with *q* = 4*π*sin(θ)/*λ*, where *λ* is the wavelength of the X-ray beam and θ is the half scattering-angle. The dimer data collected using the sample changer was inspected for radiation damage and averaged and background subtracted using primus in the ATSAS suite (Franke et al, 2017). SEC-SAXS data was collected with an in-line 24 ml Superdex 200 increase column operated at a flow rate of 0.5 ml/min. The trimer and tetramer data from SEC-SAXS were reduced using the chromixs tool from the ATSAS suite (SEC curves are presented in S2F-G). Water was used as reference to convert data to absolute scale units of cm^-1^ (32). Initial Guinier analysis to verify that no larger aggregates were present (data not shown) and indirect Fourier transformation to determine the *p(r)* functions were performed using the BayesApp software (33) available through the “GenApp.Rocks” server maintained by Emre Brookes at University of Texas. For plotting of the scattering data, the ~800 data points were logarithmically rebinned into ~180 points with better high-*q* statistics. Also, the data are only plotted in the fitted range out to about 0.25 Å^-1^. Central model independent parameters of the SAXS analysis are presented in Figure S2A.

Rigid body modelling of FP monomer and oligomers was performed in CORAL (34). The FP eye formed by the TB domain, TSR1, TSR4, TSR5 and TSR6 were used as a single rigid body. TSR2 and TSR3 from the crystal structure of FPc (entry 6RUS) formed two additional bodies that were linked to each other and the eye with distance restraints. In addition, an intact Asn linked complex glycan (entry 3RY6) formed a fourth rigid body linked to FP Asn428, whereas mannosyl groups linked to tryptophans and fucose-glucose disaccharides linked to serine and threonine were included in the same rigid body as the TSR domain they form a covalent bond with. The starting models were constructed such that two residues to be connected across the rigid bodies were in proximity. Starting models of FP oligomers were generated by C_n_ symmetry and oriented with their rotation axis along the z-axis. For each FP system, six different eyes derived from entries 6S08, 6S0A, 6S0B, 6RUS, 6SEJ, 6RUR (2,3) were first evaluated with a consistent set of distance restraints to identify the optimal eye for the final rigid body refinements. The α angle in Figure 3E-F was calculated as the angle between a vector connecting the Cα atoms of FP residues Ala402 and Ser345 and the shortest principal axis of the SAXS rigid body models. If the average α was <90, the model was flipped to yield an average α>90. Ser345 and Ala402 were chosen for definition of the FP eye vector, as their difference vector is in the plane of the FP eye and roughly parallel to the long axis of the C3b in C3bBbFP complex (2). Scattering data together with an example of output model and fit to the experimental data are deposited in the SASBDB for the FP dimer, trimer and tetramer. Scattering data for the FP E244K monomer is available as SASBDB entry SASDB69.

### Modeling and MD simulations of the FP E244K monomer

The FP1 monomer obtained by CORAL rigid-body modeling was used as the template to construct an initial atomistic model of FP E244K with Modeller9.18 (35) in which missing residues in loops connecting domains were added and the disulfide bond Cys132-Cys170 was established. Glycosylations were added including the Asn linked-glycan at Asn428, O-linked Glucoseβ1-3Fucose at Thr92, Thr151, Ser208 and Thr272, and C-mannosylations at Trp83, Trp86, Trp139, Trp142, Trp145, Trp196, Trp199, Trp202, Trp260, Trp263, Trp321, Trp324, Trp382, Trp385, and Trp388. The glycan at Asn428 was modeled as a complex glycan. FP1 E244K was placed into a periodic cubic box with sides of 14.6 nm solvated with TIP3P water molecules containing Na^+^ and Cl^-^ ions at 0.15 M, resulting in ~300,000 atoms in total. The CHARMM36m force field (36) was used for the protein. Force field parameters for N- and O-linked glycans were generated using the Glycan Modeler module in the CHARMM-GUI web interface (37). Force field parameters for C-Mannosyl Trp were obtained from Shcherbakova et al. (38). Neighbor searching was performed every 20 steps. The PME algorithm was used for electrostatic interactions with a cut-off of 1.2 nm. A reciprocal grid of 128×128×128 cells was used with 4th order B-spline interpolation. A single cut-off of 1.2 nm was used for Van der Waals interactions. MD simulations were performed using Gromacs 2019.4 or 2019.5 (39). The temperature and pressure were kept constant at 300 K using the Nose-Hoover thermostat and at 1.0 bar using the Parrinello-Rahman barostat with a time constant of 5 ps and a frequency of 20 for coupling the pressure, respectively. Two independent MD simulations (one microsecond for each) were performed to collect the conformational ensemble. These sampled conformations were used for further ensemble refinement using the Bayesian Maximum Entropy (BME) method guided by experimental SAXS data as described (40–42). By tuning the regularization parameter in the BME reweighting algorithm, we adjusted the conformational weights in variant degrees to improve the fitting with the experimental SAXS data for FP E244K. VMD and PyMol were used for visualization of the conformational ensemble and movie preparation. The theoretical SAXS curve for each frame was back calculated using Crysol3 (43).

### Single particle negative stain EM data acquisition and analysis

All samples were purified on a 24 ml Superdex 200 increase size exclusion column equilibrated in 20 mM HEPES pH 7.5, 150 mM NaCl and subsequently adsorbed to glow discharged carbon coated copper grids, washed with deionized water and stained with 2 % (w/v) uranyl formate. Images were acquired with a FEI Tecnai G2 Spirit transmission microscope at 120 kV, a nominal magnification of 67.000x and a defocus ranging from 0.7 to 1.7 μm. Automated image acquisition was performed using leginon (44). For the FP1 monomer and its hFPNb1 complex, CTF estimation and subsequent particle picking and extraction was carried out with cisTEM (45). For the remaining samples, CTF estimation, manual particle picking and extraction were performed with RELION (46). Initial 2D classes were generated and used to set up template-based particle picking. For all samples, 2D classification was performed in RELION.

### FPc and FP exchange assay

The stability of FPhtΔ3 under acidic conditions was evaluated by SEC on a 24 mL Superdex200 column (GE Healthcare) equilibrated in 20 mM HEPES, 150 mM NaCl pH 7.5 or in 100 mM glycine pH 2.3. Samples of 100 μL FPhtΔ3 at 1 mg/mL were injected and eluted at 0.5 mL/min at room temperature. Fractions were analyzed by SDS-PAGE followed by silver staining using the SilverQuest silver staining kit (Thermo Fisher). Fractions containing 0.1 M glycine pH 2.3 were neutralized with 100 μL 2 M Tris pH 8.5 before SDS-PAGE analysis. Five μg of FPc or dimeric FP were mixed with 5 μg of the his-tagged FPhtΔ3 in 100 μL of 20 mM HEPES, 150 mM NaCl pH 7.5 and were incubated on ice for 5 min or at 37 °C for 5 min, 30 min, 2 h, 6 h, 12 h, 24 h or 7 days. As a positive control, 5 μg of FPc or FP were mixed with 5 μg of FPhtΔ3 in 100 μL 20 mM HEPES, 150 mM NaCl pH 7.5 and subsequently acidified by adding 100 μL 0.1 M Glycine pH 2.3 to cause oligomer dissociation (5). The sample was incubated for 5 min at room temperature before 20 μL of 2M Tris pH 8.5 was added for neutralization. Pull downs were performed on all samples using 50 μL of Ni-NTA beads. The beads were transferred to a 1 mL spin columns (Bio-Rad) and equilibrated in 100 mM HEPES, 0.5 M NaCl, 30 mM imidazole pH 7.5. The samples were then transferred to the columns and incubated for 2 min, followed by a 30 s centrifugation step at 70 g. The beads were washed 5 times with 500 μL of 100 mM HEPES, 0.5 M NaCl, 30 mM imidazole pH 7.5 before bound protein was eluted with 80 μL of 100 mM HEPES, 0.5 M NaCl, 400 mM imidazole pH 7.5. The eluates were retrieved from the columns by a 30 s centrifugation step at 70 g. The eluates were re-applied to the column and the centrifugation step was repeated. The samples were analyzed under non-reducing conditions on a 12 % SDS-PAGE gel (GenScript) using the SilverQuest silver staining kit (Invitrogen). The oligomeric FP used in this assay was approximately 90 % dimer and 10 % trimer as judged by SEC analysis performed on 24 mL Superdex200 increase column (Fig. S1G). A SEC standard (Bio-Rad) was used for comparison.

### Bio-layer interferometry assays

Bio-layer interferometry experiments were performed on an Octet Red96 (ForteBio) at 30 °C and shaking at 1,000 RPM. Binding of hFPNb1:FPc complexes were tested on streptavidin sensor tips (SA, ForteBio) equilibrated in assay buffer (PBS supplemented with 1 mg/mL BSA and 0.05 % Tween 20) and coated with biotinylated C3b at 16 μg/mL for 10 min. FPc or hFPNb1:FPc complexes were prepared by size exclusion chromatography and diluted to 50 μg FPc/mL in assay buffer. Association was monitored for 120 s followed by 120 s dissociation in assay buffer.

### Erythrocyte lysis assay for AP activity

Rabbit erythrocytes (Er) in 11.38 mM D-Glucose, 2.72 mM mono basic sodium citrate, 2.19 mM citric acid, 7.19 mM NaCl (Alsever’s Solution, Statens Seruminstitut) were washed and re-suspended in AP assay buffer (5 mM barbital, 145 mM NaCl, 10 mM EGTA, 5 mM MgCl2, pH 7.4, with 0.1 % (w/v) gelatin) to obtain a 6 % (v/v) suspension. Samples of 20 μL human FP-deficient serum diluted in assay buffer and supplemented with WT FP, FPc or hFPNb1-FPc complexes were prepared separately and then transferred to a V-shaped bottom 96-well microtiter plate (Nunc) in duplicates. The WT FP used for this assay was approximately 50 % dimer and 50 % trimer as judged by SEC analysis performed on a 24 mL Superdex200 increase (Fig. S1F). Ten micro liter of the Er-suspension was then transferred to the assay plate. The plate was mixed well and incubated for 2 h at 37 °C, and shaken every 30 min. Hemolysis was stopped by adding 40 μL ice-cold 0.9 % NaCl, 5 mM EDTA to each well. The plate was centrifuged at 90 *g* for 10 min, and 50 μL of each supernatant were subsequently transferred to a flat-bottom microtiter well plate (Nunc). Hemolysis was then determined from the A405 measured on a Victor3 plate reader (PerkinElmer). Results are expressed relative to total hemolysis (obtained with water alone) and to background hemolysis (EDTA sample).

---

**Figure S1.**
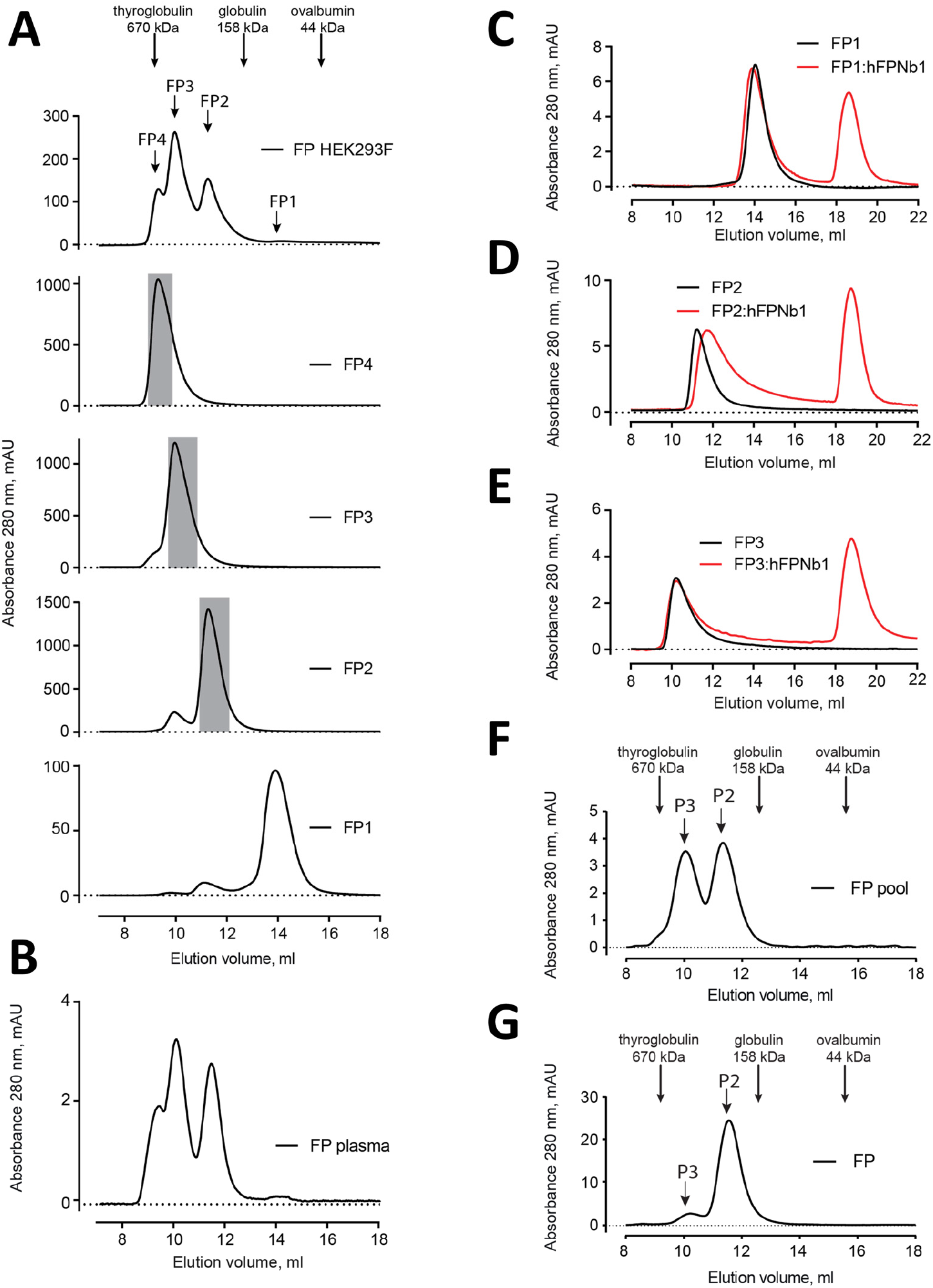
Elution profiles of recombinant FP. A) Top, SEC analysis of initial pool of recombinant FP obtained by His_6_ based affinity chromatography. Below is presented SEC profiles for the final purification step of recombinant FP1, FP2, FP3 and FP4 obtained from HEK293F cells used for SAXS data collection and negative stain EM (greyed areas). B) SEC profile of commercial plasma FP for comparison. C-E) SEC analysis of FP1, FP2, and FP3 with or without hFPNb1 used for nsEM in figures 1 and 2. F-G) SEC analysis of the FP2/FP3 pool and a pool dominated by FP2 used for the functional assays in figure 4 and exchange experiments in figure S4, respectively.

**Figure S2.**
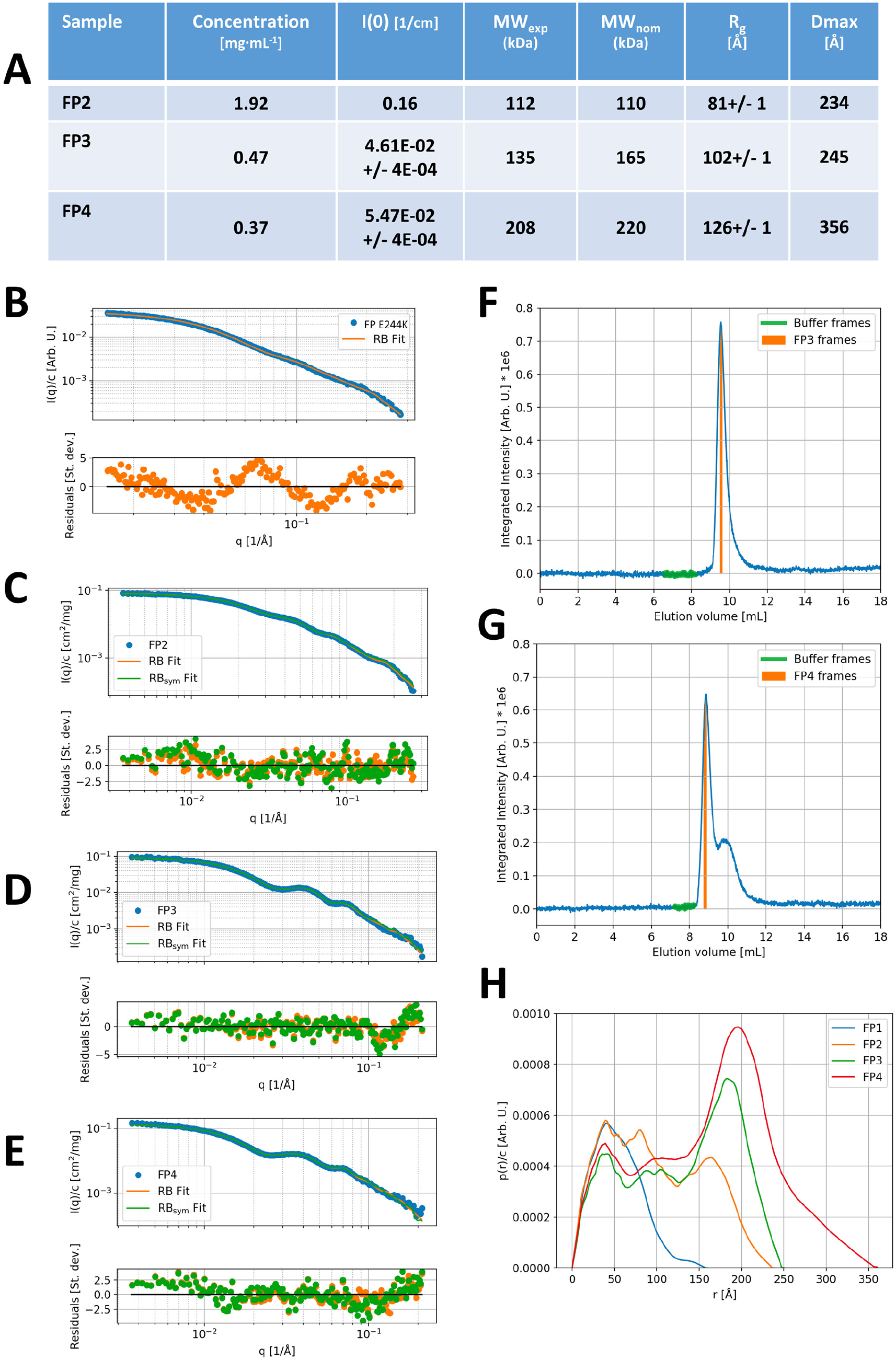
SAXS data. A) Model independent parameters of the SAXS analysis. Forward scattering intensity, *I(0)*, in absolute units of cm^-1^. MW_exp_: Average MW of the oligomer as derived from the forward scattering and the protein concentration. The uncertainty of the MW determination is expected to be around 10% for the FP2 which was measured using the sample changer setup and around 20% for the SEC-SAXS based setup. When taking this into account, the experimental MW-values compare well with their nominal values, MW_nom_, calculated using a monomer MW of 55 kDa. *R_g_*: Radius of gyration and *D_max_*: Maximal distance of the *p(r)*. The *I(0), R_g_* and *D_max_* are derived from the Indirect Fourier Transformation as described in the materials and methods section. B-E) Experimental SAXS data (blue points), fit of rigid body (RB) model (orange full line) and fit of C_n_ symmetric model with n=2 (FP2), 3 (FP3) and 4 (FP4), respectively, in panels C, D and E (green full line). Corresponding residual plots are presented underneath each SAXS plot. F-G) Integrated SAXS intensity of the SEC-SAXS data obtained for FP3 and FP4. The frames used for the foreground (FP3/4 frames) and buffer frames selected for the background are indicated on the plots. H) Direct comparison of the *p(r)* functions obtained from the SAXS data on FP1 E244K, FP2, FP3 and FP4.

**Figure S3.**
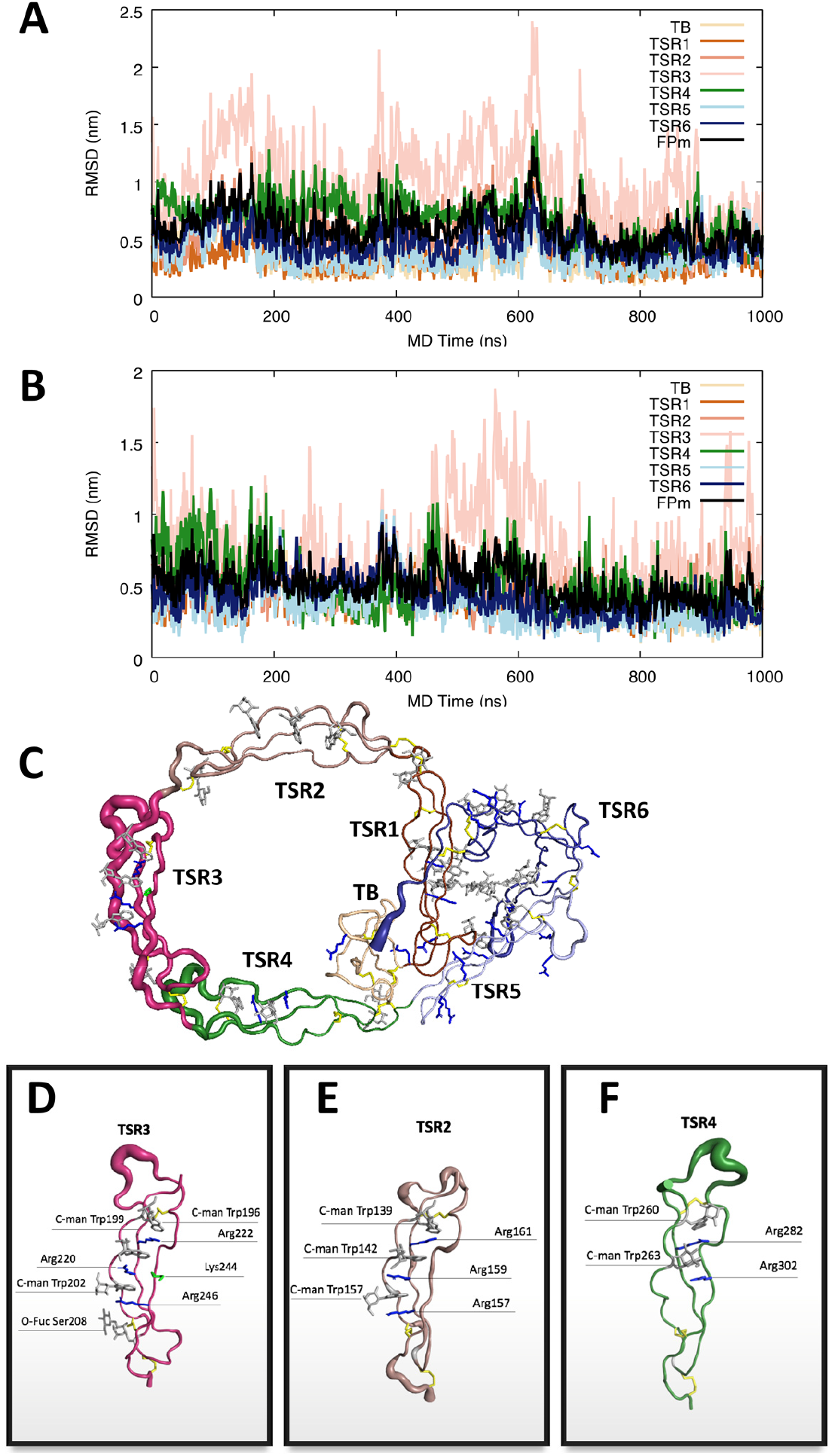
Molecular dynamics simulation of the FP1 E244K. A-B) The root mean square deviation (RMSD) of each domain and the whole protein with the trajectories aligned to the initial FP1 E244K model in two independent one μs MD simulations. Both simulations commonly revealed significant dynamics of TSR3. C) The RMS fluctuations within the entire FP E244K with the trajectory aligned to the initial model illustrated by cartoon representations in which the thickness of the tube is scaled by the corresponding RMS fluctuation values per residue. D-F) Close-up of the RMS fluctuations of TSR3, TSR2 and TSR4 with the trajectories aligned to the corresponding domain, indicating that these thrombospondin repeats have limited conformational flexibility in their overall structure. Importantly, the mutation of glutamate 244 to lysine in TSR3 does not cause breakdown of the central Trp-Arg stack.

**Figure S4.**
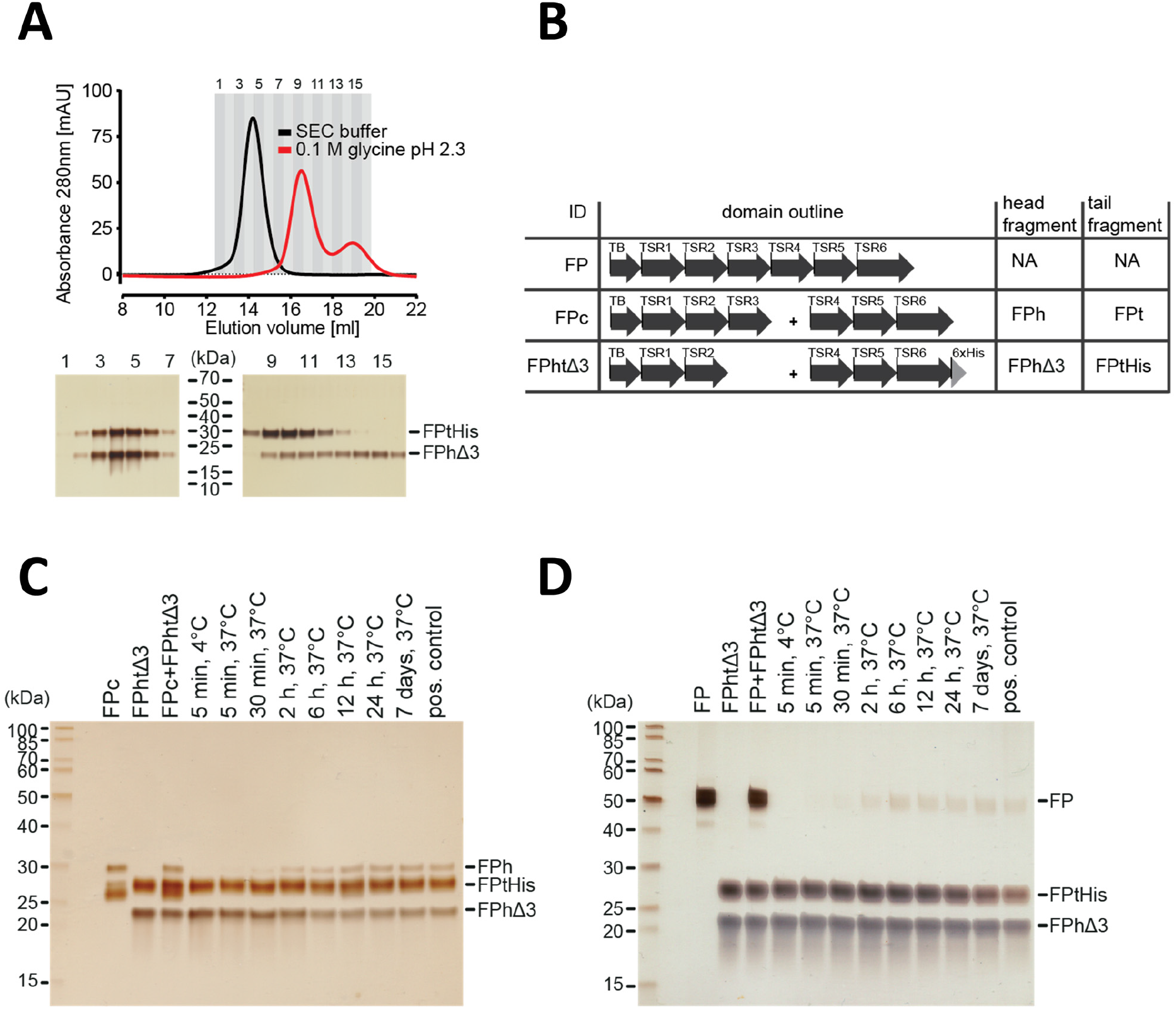
FP oligomerization interfaces can open up to enable exchange monomer. A) SEC analysis of FPhtΔ3 in 20 mM HEPES, 150 mM NaCl, pH 7.5 (black) or 100 mM glycine pH 2.3 (red), demonstrating that the two chains of FPhtΔ3 dissociate at low pH (top). Fractions from the SEC experiments were analyzed by SDS-PAGE and silver staining showing that at neutral pH the two fragments of FPhtΔ3 co-elute (bottom, left) whereas at low pH FPtHis (25 kDa) elute first and FPhΔ3 (18 kDa) elute last (bottom, right). B) Overview of FP constructs used in panels C-D. C) A silver-stained SDS-PAGE gel following the exchange of chains between FPc and FPhtΔ3. The exchange was monitored by the appearance of the FPh chain (25 kDa) from FPc and the disappearance of the FPhΔ3 chain, after pull-down on Ni-NTA beads using the His-tag on the C-terminus of FPtHis. D) A silver-stained gel from SDS-PAGE demonstrating the exchange of FP chains between FP2 and FPhtΔ3. The exchange was monitored by the appearance of the full length FP (50 kDa) after pull-down on Ni-NTA beads using the C-terminal His-tag of FPtHis from FPhtΔ3.

**Movie S1. Molecular dynamics simulation of FP E244K.** Domains and glycans are colored as in figure 1F. All frames in a 1 μs simulation are displayed as thin lines in the background. Notice how TSR3 is able to rotate substantially around the two hinges connecting it to TSR2 and TSR4 facilitated by flexibility of the N-terminal regions in TSR3 and TSR4. The rest of the molecule exhibits only limited dynamics.

**Movie S2. Close-up on TSR3 during the MD simulation of FP E244K.** The thrombospondin repeat is shown in the orientation and coloring used in Figure S3D. Notice how the central Trp-Arg stack remains stable despite the presence of a lysine at position 244. In the structure of wild type FPc, the glutamate at position 244 engages the side chain of Arg220 in an electrostatic interaction and thereby stabilizes the arginine conformation.

